# Transcriptome Analysis Reveals Skin Lipid Metabolism Related to Wool Diameter in Sheep

**DOI:** 10.1101/051359

**Authors:** Shaoyin Fu, YunXia Qi, Xiaolong He, Lai Da, biao Wang, rigele Te, jianghong Wu, ding Yang, yongbin Liu, wengguang Zhang

**Author notes:** Co-first author (Shaoyin Fu, YunXia Qi).

## Abstract

Wool is one of the most important animal fibers in the textile industry and the diameter directly affects its economic value. However, the molecular mechanisms underlying the wool diameter have not been fully elucidated. In the present study, high-throughput RNA-Seq technology was employed to explore the skin transcriptome using 3 sheep with fine wool (fiber diameter, FD<21.0μm) and 3 sheep with coarse wool (fiber diameter, FD>27.0μm). In total, 28,607,228 bp clean reads were obtained, and 78.88%+/-3.84% was uniquely aligned to the reference genome across the six samples. In total, 19,914 mRNA transcripts were expressed (FPKM>0) in the six skin samples, among which there were certain well-known genes affecting the skin hair cycle, such as KRTAP7-1, KRT14, Wnt10b, Wnt2b, β-catenin, and FGF5. Furthermore, 467 expressed genes were significantly differentially expressed between the two groups, including 21 genes up-regulated and 446 genes down-regulated in the sheep with the smaller fiber diameter. To verify the results, 13 differentially expressed genes were randomly selected to validate the expression patterns using qRT-PCR, and the correlation between the mRNA expression level from qRT-PCR and RNA-Seq data was 0.999 ( P<0.05). These differentially expressed genes were particularly enriched in GO processes related to lipid metabolism, skin development, differentiation, and immune function (P<0.05). The biological processes were involved in collagen catabolism, negative regulation of macromolecule metabolism, steroid hormone stimulation and lipid metabolism. A significant KEGG pathway involving the “metabolism of lipids and lipoproteins” was also enriched. This study revealed that the lipid metabolism might constitute one of the major factors related to wool diameter.

## Introduction

Sheep (*Ovis aries*) is a predominant domestic animal providing not only meat and milk but also wool. Wool is a distinguishing feature of sheep compared to other farm animals, and its biology has been the focus of much research. Wool is one of the earliest natural fibers used in the textile industries. The wool fiber is soft and elastic, and its products possess the advantage of being natural, having strong hygroscopicity, providing warmth and comfort, etc. The key traits contributing to the economic value of wool include fiber diameter, density, strength and length which are determined by both genetics[1,2] and the environment[3]. Understanding the genetic principles of wool traits would be helpful for promoting sheep breeding and also for elucidating the mechanism of hair development in humans.

Over the past few decades, progress has been made on the study of wool quality using genetic technologies. In the beginning, these studies mainly focused on the wool biology and the quantitative trait loci associated with wool economic traits, and some major genes have been validated. For example, KRTAP6 and KRTAP8, located in Chromosome 1[4], control wool diameter andagouti is a key locus affecting wool color. The N-type gene, also named “halo-hair 1” (HH1) gene, is another important gene controlling wool quality, as mutations in this gene cause extreme hairiness (or medullation), resulting in the production of fibers that are ideal for carpet wool production[5,6].

With rapid development of molecular biological techniques, especially the Next-generation sequencing, large-scale gene expression detection becomes possible. Fan *et al*. [2] examined the skin gene expression profiles associated with coat color in sheep using RNA sequencing (RNA-Seq). Kang *et al*. [3] studied characteristics of curly fleece utilizing transcriptome data. Yue *et al*. [7,8] performed *de novo* transcriptome sequencing of sheep skin and found fiber diameter was related to a few lipoic acid differentially expressed genes and antisense transcripts. However, no other studies have examined global gene expression in relation to wool fiber diameter. To gain better understanding of molecular mechanisms controlling wool fiber diameter, global gene expression profiles in skin of sheep with coarse wool versus fine wool were explored. The results showed that wool fiber diameter is associated with lipid metabolism and provided valuable information for future studies.

## Materials and Methods

### Experimental Animals and Sample Collection

The experimental animals are Erdos Fine Wool sheep from Inner Mongolia Autonomous Region, China. Three hundred and sixty unrelated female ewes, that were 3 years old and raised in the same conditions, were randomly selected from a sheep farm. The diameter of wool from each animal was measured and recorded. According to the fineness of the wool, 3 animals of fine wool (fiber diameter, FD<21.0μm, FW) and coarse wool (fiber diameter, FD>27.0μm, CW) were sampled. A piece of skin (approximately 0.5cm^2^) on the body side was cut off and frozen immediately in liquid nitrogen and stored at −80°C until subsequent use.

The experiment was conducted following the Guidelines on Ethical Treatment Animals (2006 No.398), set by the Ministry of Science and Technology, China. The sampling procedures complied with the Animal Ethics Committee at Inner Mongolia Academy of Agricultural & Animal Husbandry Sciences.

### RNA Extraction, cDNA Library Construction and Illumina Sequencing

RNA from the two groups was extracted using Trizol reagent (TaKaRa) according to the manufacturer’s instructions, and the purity and degradation were determined on 1% agarose gels. DNA was removed from the RNA extracts by incubation with RNase-free DNase for 30 min at 37°C.

Poly(A) mRNA was isolated from total RNA using oligo(dT) magnetic beads (Illumina). Fragmentation buffer was added to disrupt the purified mRNA into short fragments. Using these short fragments as templates, first-strand cDNA synthesis was performed using random hexamer primers and reverse transcriptase (Illumina). Second-strand cDNA was synthesized using RNase H (Illumina), DNA polymerase I (Illumina), dNTPs and buffer. These cDNA fragments were subjected to end repair process and ligation of adapters. These products were purified and enriched with PCR to create the finally cDNA library. The library preparations were sequenced on an Hiseq 2000 platform and 100 bp paired-end reads were generated.

### Sequence preprocessing and functional annotation

In order to obtain clean data, all sequenced raw data were processed including removing the adapter, filtering the low quality reads and the proportion of N more than 10% using an in-house Perl script. Then, all filtered data were mapped to the sheep genome (V 3.1) using Tophat2, with no discordant and mixed. Unique mapped reads were kept to estimate the gene abundances in the downstream analysis. Reference guided transcriptome assembly, which compensates incompletely assembled transcripts, was performed by Cufflinks with bias correction for each sample, and then merged into a single unified transcript catalog using Cuffmerge discarding isoforms with abundance below 0.1 in order to remove the low-quality transcripts. As well, differently expressed genes were estimated by Cuffdiff using FPKM [9] (Reads Per Kilobase per Million mapped reads) based on P-value<0.05.

### Enrichment analysis

To explore the functional annotation and pathway enrichment of those significantly different genes interactive between the fine and coarse wool, the on-line analysis tool GSEA (GSEA v2.1.0 :software.broadinstitute.org/gsea/msigdb/annotate) was used to determine the enriched Gene Ontology (GO) terms and Kyoto Encyclopedia of Genes and Genomes ( KEGG) (P-value<0.05).

### Quantitative real time PCR (RT-PCR) validation

Three animals with fine wool (fiber diameter, FD<21.0μm) and three animals with coarse wool (fiber diameter, FD>27.0μm) were selected. A piece of skin (approximately 0.5cm^2^) on the body side was cut off and frozen immediately in liquid nitrogen for the subsequent qRT-PCR analysis. Total RNA was extracted using Trizol (TaKaRa) following the manufacturer’s protocols. The total RNA obtained was re-suspended in nuclease-free water and the concentration was measured using Nanodrop (Thermo Scientific Nanodrop 2000). Approximately 0.5 μg of total RNA was used as template to synthesize the first-strand cDNA using a PrimerScript RT reagent Kit (TaKaRa) following the manufacturer’s protocols. The resultant cDNA was diluted to 0.1 μg/μl for further analysis in the qRT-PCR (Bio-Rad) using a SYBR Green Realtime PCR Master Mix (TaKaRa). The GAPDH gene was chosen as internal reference to eliminate sample-to-sample variation. The relative gene expression levels were calculated using the 2^−ΔCt^ method. The differential expression genes between fine and coarse wool skin samples were analyzed by GLM using SAS software 9.0.

## Results

### Skin transcriptome profiling of differential diameter

The cDNA libraries of six skin samples from two groups of fine wool sheep with different fiber diameters (3 samples for FD<21.0μm and 3 samples for FD>27.0μm) were sequenced. In total, 35,884,100-55,926,820 paired-end reads of 100 bp in length were obtained per sample. As a result, the total read length was 28.6 gigabases (Gb) for the six samples. Alignment of the sequence reads against the Ovis aries 3.1 reference genome sequence (www.ncbi.nlm.nih.gov/genome/?term=sheep) yielded 78.88%+/-3.84% of uniquely aligned reads across the six samples, of which 63-67% was in annotated exons, 10-11% was located in introns, and the remaining 22-27% was assigned to intergenic regions. Unmapped or multi-position matched reads (7.6-8.5%) were excluded from further analyses. Consequently, 19914 mRNA transcripts were detected as expressed (FPKM>0) in the six skin samples (Additional file1:Table S1). Among these, there were several well-known genes affecting skin cycle (e.g., *KRTAP7-1, KRT14, Wnt10b, Wnt2b, β-catenin*, and *FGF5*).

### Differential gene expression between two groups for fine wool and coarse wool

Using Cuffdiff methods [9], the differential gene expression profile between the Erdos fine wool sheep with large and small fiber diameters were examined. In total, 467 expressed genes were detected as significantly different on the basis of the threshold value (*P<0.05*, |log_2_Ratio|>1.4). The expression levels of 21 of the 467 genes were upregulated in the sheep with small fiber diameter; the other 446 genes showed higher expression in the sheep with coarse wool (Figure 1 and Table S2).

**Figure 1.**
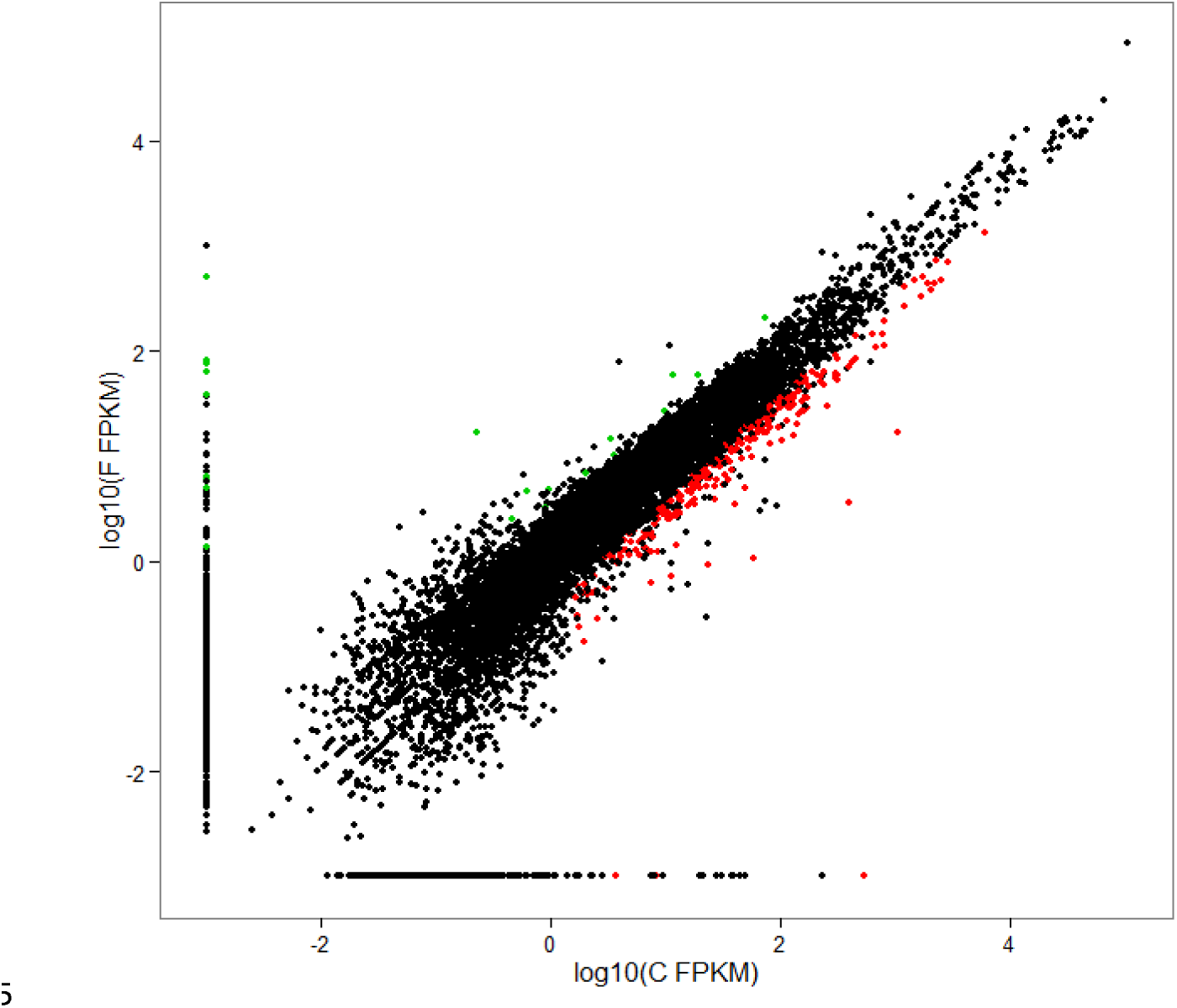
Scatter plot displaying differential expressed genes between the skin samples of two groups of sheep with different fiber diameters. The red dots represent up regulated genes in coarse wool group, the green dots represent up regulated genes in fine wool group. In total, 467 genes were identified as differentially expressed (P-value b<0.05 & |log2Ratio|>1.4) between F (F means Fine wool group) and C (C means coarse wool group). FPKM: Fragment Per Kilobase of exon model per Million mapped reads.

The reliability of the RNA-Seq data and the sampling accuracy of the skin tissue used in this study were confirmed by qRT-PCR of 13 randomly selected genes. The 13 differentially expressed genes included *GNPAT, UGCG, PLD2, ACOT8, FABP4, PLA2G3, LPIN2, SMPD2, HSD11B1, FA B P 6, CHKA, SLC25A20* and *NCOR1*. For these selected 13 genes, the correlation between the mRNA expression level from qRT-PCR and RNA-Seq were high (correlation coefficient = 0.999, *P*<0.05), confirming the high reproducibility of RNA-Seq data in this study (Figure 2 and Table S3).

**Figure 2.**
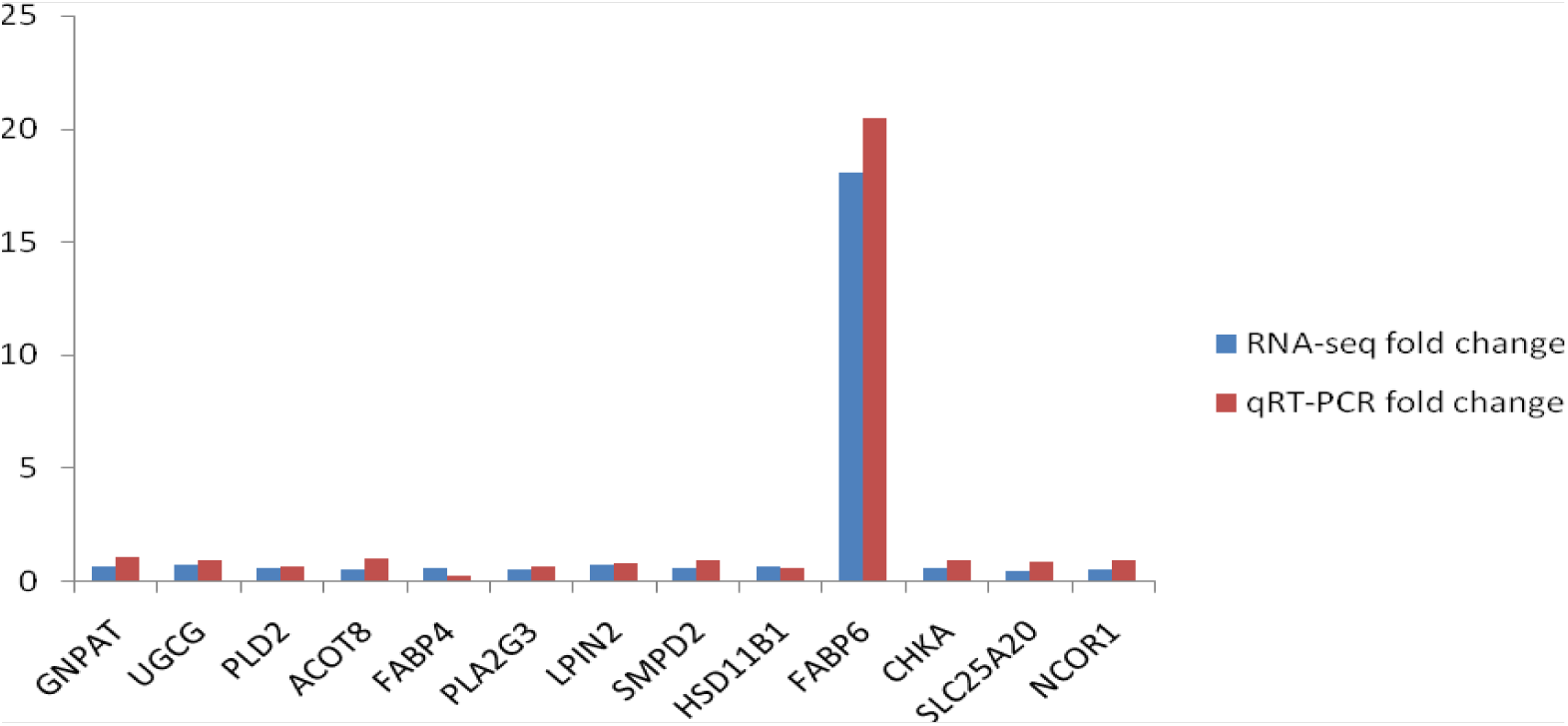
qRT-PCR validation of differentially expressed genes. For the 13 randomly selected differentially expressed genes, fold changes of FW/CW determined from the relative Ct values using the 2-^Δ^Ct method in qRT-PCR were compared to those detected by FPKM of FW/CW in RNA-Seq. All Ct values were normalized to GAPDH and replicates (n = 3) of each sample were run.

### Gene Ontology enrichment and pathway analysis

To further investigate the functional association of the 467 differential expressed genes, gene ontology (GO) analysis using the on-line analysis tool GSEA (GSEAv2.1.0 :software.broadinstitute.org/gsea/msigdb/annotate) was performed. Several significant GO categories were enriched (*P<0.05*), including the GO processes related to lipid metabolism, skin development, differentiation, and immune function. Such biological processes were involved in collagen catabolism, negative regulation of macromolecule metabolism, steroid hormone stimulation and lipid metabolism (Table 2).

**Table 2.** **GO analysis of 467 differentially expressed genes**

| Go ID | Go term | No.of differentially | P-value |
| --- | --- | --- | --- |

|  |  | expressed genes |  |
| --- | --- | --- | --- |
| GO:0030574 | collagen catabolic process | 4 | 4.84E-03 |
| GO:0010605 | negative regulation of macromolecule metabolic process | 24 | 0.004948 |
| GO:0048545 | response to steroid hormone stimulus | 10 | 0.006725 |
| GO:0010558 | negative regulation of macromolecule biosynthetic process | 19 | 0.007727 |
| GO:0031324 | negative regulation of cellular metabolic process | 23 | 0.007983 |
| GO:0048523 | negative regulation of cellular process | 43 | 0.008742 |
| GO:0009892 | negative regulation of metabolic process | 24 | 0.010027 |
| GO:0044243 | multicellular organismal catabolic process | 4 | 0.010223 |
| GO:0009890 | negative regulation of biosynthetic process | 19 | 0.012006 |
| GO:0032963 | collagen metabolic process | 4 | 0.012555 |
| GO:0006730 | one-carbon metabolic process | 7 | 0.013863 |
| GO:0006629 | lipid metabolic process | 24 | 0.015868 |
| GO:0044259 | multicellular organismal macromolecule metabolic process | 4 | 0.01658 |
| GO:0031327 | negative regulation of cellular biosynthetic process | 18 | 0.019938 |
| GO:0048519 | negative regulation of biological process | 44 | 0.022378 |
| GO:0010551 | regulation of specific transcription from RNA polymerase II promoter | 6 | 0.024569 |
| GO:0044236 | multicellular organismal metabolic process | 4 | 0.026557 |
| GO:0009725 | response to hormone stimulus | 13 | 0.028269 |
| GO:0008152 | metabolic process | 150 | 0.028768 |
| GO:0032583 | regulation of gene-specific transcription | 7 | 0.030424 |
| GO:0044238 | primary metabolic process | 137 | 0.032539 |
| GO:0010553 | negative regulation of specific transcription from RNA polymerase II promoter | 4 | 0.034637 |
| GO:0043627 | response to estrogen stimulus | 6 | 0.037124 |
| GO:0016070 | RNA metabolic process | 25 | 0.039369 |
| GO:0005739 | mitochondrion | 31 | 0.015305 |
| GO:0005622 | intracellular | 218 | 0.018333 |
| GO:0005737 | cytoplasm | 152 | 0.021306 |
| GO:0043227 | membrane-bounded organelle | 162 | 0.038967 |
| GO:0044424 | intracellular part | 209 | 0.039617 |
| GO:0016607 | nuclear speck | 6 | 0.040209 |
| GO:0043231 | intracellular membrane-bounded organelle | 161 | 0.048547 |
| GO:0016787 | hydrolase activity | 72 | 1.39E-06 |
| GO:0003824 | catalytic activity | 125 | 6.91E-05 |
| GO:0016462 | pyrophosphatase activity | 27 | 0.001184 |
| GO:0016818 | hydrolase activity, acting on acid anhydrides, in phosphorus-containing anhydrides | 27 | 0.001266 |
| GO:0004175 | endopeptidase activity | 17 | 0.001274 |
| GO:0016817 | hydrolase activity, acting on acid anhydrides | 27 | 0.001357 |
| GO:0017111 | nucleoside-triphosphatase activity | 26 | 0.001477 |
| GO:0008233 | peptidase activity | 21 | 0.003694 |
| GO:0070011 | peptidase activity, acting on L-amino acid peptides | 20 | 0.004918 |
| GO:0004252 | serine-type endopeptidase activity | 9 | 0.006701 |
| GO:0004386 | helicase activity | 8 | 0.013278 |
| GO:0008236 | serine-type peptidase activity | 9 | 0.01523 |
| GO:0017076 | purine nucleotide binding | 48 | 0.016065 |
| GO:0017171 | serine hydrolase activity | 9 | 0.016177 |
| GO:0032553 | ribonucleotide binding | 46 | 0.018382 |
| GO:0032555 | purine ribonucleotide binding | 46 | 0.018382 |
| GO:0003723 | RNA binding | 22 | 0.019358 |
| GO:0000166 | nucleotide binding | 54 | 0.020723 |
| GO:0005524 | ATP binding | 38 | 0.023655 |
| GO:0030554 | adenyl nucleotide binding | 40 | 0.024429 |
| GO:0032559 | adenyl ribonucleotide binding | 38 | 0.028346 |
| GO:0001883 | purine nucleoside binding | 40 | 0.030096 |
| GO:0004089 | carbonate dehydratase activity | 3 | 0.032682 |
| GO:0001882 | nucleoside binding | 40 | 0.033016 |
| GO:0016836 | hydro-lyase activity | 4 | 0.044234 |
| GO:0003714 | transcription corepressor activity | 7 | 0.046929 |
| GO:0005044 | scavenger receptor activity | 4 | 0.049408 |

In addition, a pathway analysis of the 467 differentially expressed genes, using GSEAv2.1.0, was also performed. The KEGG pathway involving “metabolism of lipids and lipoproteins” was significantly enriched. In total, 16 genes (*GNPAT, LPIN2, CHKA, PLD2, PLA2G3, SMPD2, UGCG, PIP4K2A, ACOT8, SLC25A17, NCOR1, FABP6, HSD11B1, STAR, FABP4* and *SLC25A20*) were found to be involved in the metabolism of lipids and lipoproteins pathway were significantly related to fiber diameter (P<0.01) (Table 3).

**Table 3.**
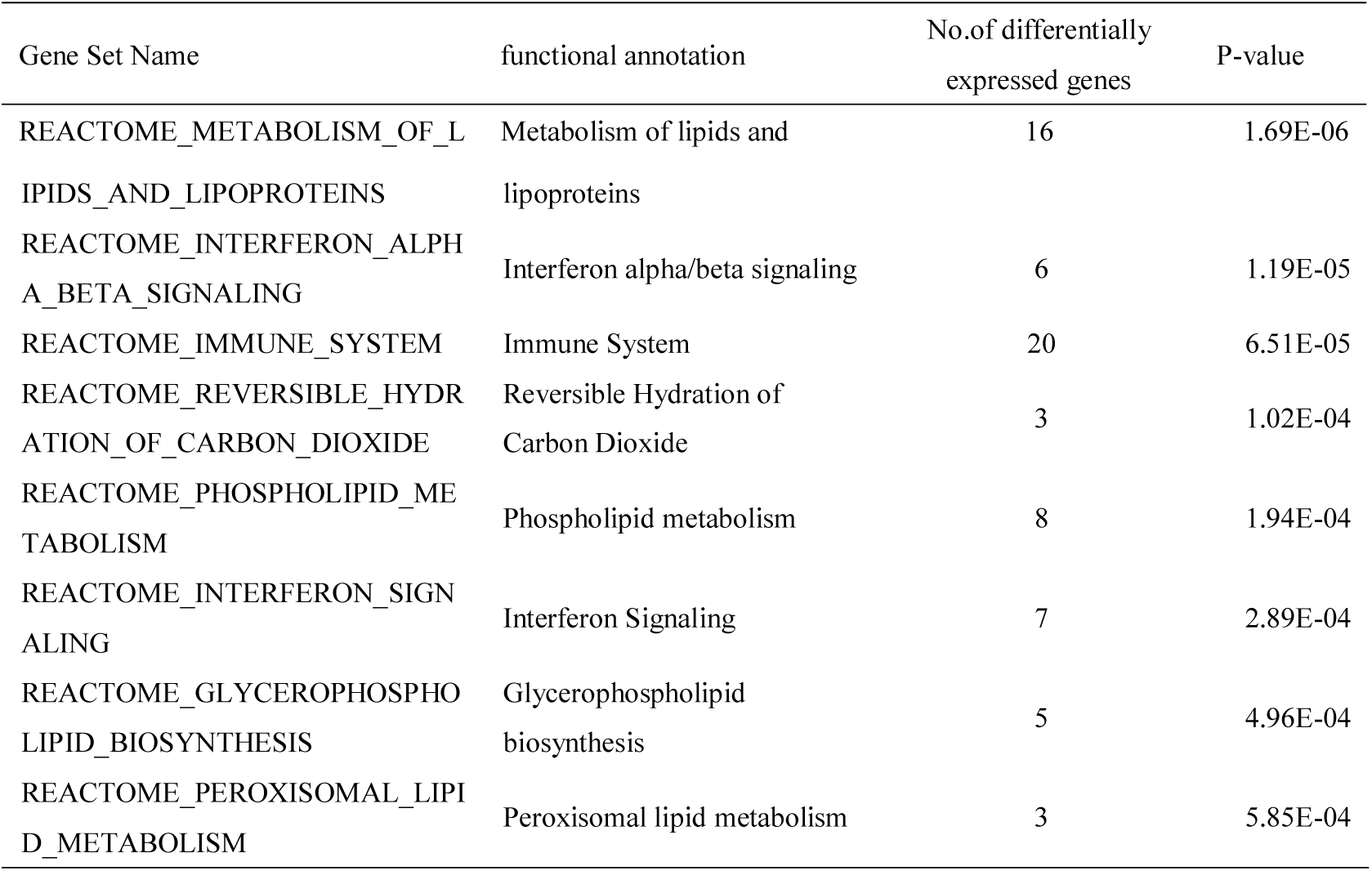
**KEGG analysis of 467 differentially expressed genes**

## Discussion

The mechanisms controlling wool/hair traits are complicated. Wool and hair fibers are composed of approximately 90% protein and 1- 9% lipid (dry weight) [10]. Thus, synthesis of wool may be linked to lipid metabolism [11]. However, studies on wool traits and regulation mechanisms have been mostly concentrated on the proteins, especially Keratin intermediate filament (KRT-IF) and Keratin-associated proteins (KAPs) [12–14]. The relationship between wool lipids and wool traits, however, has not been examined in detail. Wool lipids are designated as external and structural (or internal) based on their location in the fiber [15]. External lipids, namely wool grease (lanolin) which is secreted from the sebaceous glands attached to the wool follicles, constitutes 10-25% of the wool weight [16]. Internal lipids represent approximately 1.5% of the wool weight, consisting mainly of free fatty acids (FFA), sterols and ceramides [17]. Only a few previous studies [18–20] have reported on the wool and hair lipid composition, structural arrangement and physicochemical properties.

Recently, researchers have discovered by sheep genome and transcriptome sequencing that some genes involved in skin lipid metabolism, such as *LCE7A* and *MOGAT3*, may be related to wool synthesis [11]. Employing X-ray and molecular dynamics simulation, analysis of the effect of the internal lipids on alpha-keratin protein showed that excess internal hair lipids intercalate a dimer of keratin to disorganize the ordered keratin structure [10]. To demonstrate the important role of lipids on hair keratin structure, Cruz *et al*. (2013) removed the lipids from the simulated keratin/lipids mixture to allow the keratin to organize itself. As lipids intercalate with keratin structure, they may affect the tensile strength of the hair keratin. Moreover, higher lipid content may have the ability to interact and interfere with the structure of keratin fibers, which may influence in the texture of the hair or wool [10]. A study by Duvel *et al*. [21] showed that decreased free polar lipid concentrations and covalently bound fatty acids from the root to the tip of the hair markedly decreased tensile properties. All of these studies indicate that the lipid metabolism has close relevance to wool synthesis, and may even affect wool traits.

In the present study, the gene expression profiles of sheep skin with fine and coarse wool were compared. There were 467 differentially expressed genes detected, and a large proportion of these genes were enriched in several significant GO processes related to lipid metabolism and a number of genes were found to be involved in the metabolism of lipids and lipoproteins using KEGG pathway analysis. These findings, together with previous studies, suggest that lipid metabolism may be an important mechanism affecting wool synthesis and properties.

The phospholipase gene family encodes enzymes that hydrolyze phospholipids into fatty acids and other lipophilic molecules. This gene family is classified into four major classes, namely, phospholipase *PLA, PLB, PLC* and *PLD* [22,23], based on the types of catalytic reaction of phospholipids. The majority of coding enzymes play crucial roles in lipid metabolism, cell proliferation, muscle contraction and inflammation [24–28]. PLA2G3 (group III phospholipase A2) in mammals is a multi-domain protein with a central 150 amino acid (AA) PLA2 domain flanked by N-terminal (130 AA) and C-terminal (219 AA) extensions of unknown function [29]. It was shown recently that *PLA2G3* contributes to sperm maturation, development of atherosclerosis in mice, and mast cell maturation and function in addition to other roles [25,30,31]. *PLD2* (phospholipase D2) gene is widely expressed in various tissues, and two transcript variants encoding different isoforms have been found for this gene [32]. In addition to the common roles of the phospholipase gene family, *PLD2* has been also discovered recently to catalyze a transphosphatidylation reaction to produce a phosphatidylalcohol, a potential lipid messenger regulating keratinocyte proliferation and differentiation [33]. This protein localizes to the peripheral membrane and may be involved in cytoskeletal organization, cell cycle control, transcriptional regulation, and/or regulated secretion.

Intracellular fatty acid-binding proteins (FABPs) are members of a multigene family encoding ∼ 5-kDa proteins, which bind a hydrophobic ligand in a non-covalent, reversible manner [34,35]. These proteins are thought to have various functions including fatty acid uptake, transport, and metabolism [36–41]. Nine separate mammalian FABPs have been identified to date, and each has unique tissue-specific distribution pattern [42,43]. *FABP4* encodes the fatty acid-binding protein found in adipocytes, and is used to predict the development of the metabolic syndrome independently of pubertal status, adiposity, and insulin resistance [44,45]. The FABP6 protein is found in the ileum, ovary and adrenal gland [42,46,47], and has been hypothesized to function as a cytosolic receptor for bile acids transported by the sodium dependent action of the ileal bile-acid transporter [42,48]. In the present study, both proteins were found to be expressed in skin, indicating that FABPs may have other roles.

Lipin family proteins, including lipin1, lipin2 and lipin3, are emerging as critical regulators of lipid metabolism. Specifically, they act as phosphatidate phosphatase (PAP) enzymes required for glycerolipid biosynthesis and as transcriptional coactivators that regulate expression of lipid metabolism genes [48,49]. Each of the three lipin genes exhibits a unique pattern of tissue expression, suggesting independent physiological roles. Lipin2 (*LPIN2*) is expressed in many tissues including liver, kidney, brain, and lung [50]. Several missense mutations in *LPIN2* have been associated with psoriasis [51]. Also, homozygous and compound heterozygous mutations in human *LPIN2* lead to Majeed Syndrome characterized by recurrent multifocal inflammation of bone and skin, fever, and dyserythropoietic anemia [52–55]. These findings suggest *LPIN2* functions are crucial for normal skin metabolism.

11β-Hydroxysteroid dehydrogenase 1 (*HSD11B1*) is a member of the short chain dehydrogenase/reductase superfamily [56]. The first mammalian *HSD11B1* to be cloned was a cDNA from rat liver, and analysis indicated an 861-bp open reading frame encoding a protein of 288 amino acids. Subsequently, cDNA sequences have been published for many species including humans, which are over 30 kb in length and consist of 6 exons and 5 introns [56]. *HSD11B1* has been identified in a wide variety of tissues [56,57], including skin [58]. *HSD11B1* is an endoplasmic reticulum membrane enzyme and serves primarily to catalyze reversibly the conversion of cortisol to the inactive metabolite cortisone, and the conversion of 7-ketocholesterol to 7-beta-hydroxycholesterol [56,57,59]. Recent observations also indicated a role for *HSD11B1* in oxysterol metabolism and in bile acid homeostasis [59]. Too much cortisol can lead to Cushing’s syndrome [60]. Patients with this disease have severe skin atrophy and impaired wound healing [61]. Therefore, it is considered that topical *HSD11B1* inhibition could have a beneficial impact on the ageing skin phenotype and wound healing [56,60].

## Conclusions

This study has greatly expanded our comprehension of molecular mechanisms affecting wool properties particularly wool fineness and potentially human hair texture. Differences were found in the expressed genes suggested GO categories and pathways between the two groups with different wool fiber diameters that may be related to wool fineness. These findings provide a better understanding of wool physiology and will be useful for identification of genes associated with wool diameter.

